# Fully Automated and Noise-Robust SEEG Electrode Localization under Practical Clinical Conditions

**DOI:** 10.64898/2026.02.12.705628

**Authors:** Cheng Chen, Long Zhang, Pei Feng Tong

**Affiliations:** Peking University Changsha Institute for Computing and Digital Economy, Changsha, Hunan 410205, China; Department of Statistics and Data Science, Tsinghua University, Beijing 100084, China

**Keywords:** Epilepsy, Stereoelectroencephalography (SEEG), Contacts localization, Co-registration, Clinical automated pipeline

## Abstract

Stereoelectroencephalography (SEEG) provides direct intracranial recordings of epileptic activity with high spatial and temporal resolution and is widely regarded as the gold standard for presurgical evaluation of drug-resistant epilepsy. Accurate localization of SEEG contacts is essential for seizure onset analysis and surgical planning. However, existing localization approaches are often semi-automated, sensitive to imaging quality, and require manual intervention, limiting scalability and routine clinical deployment. We present a fully automated and noise-robust SEEG contact localization pipeline that integrates adaptive artifact suppression, data-driven thresholding, and flexible geometry-based reconstruction. The method was evaluated on six patients from two centers, including diverse resolution CT acquisitions, comprising 57 electrodes and 652 contacts in total. The proposed framework successfully reconstructed all electrodes and achieved an overall recall of 98.3% and precision of 97.1% for contact localization compared with expert labeling, with a mean deviation of 0.248 ± 0.066 mm. Processing time was approximately two minutes per subject on standard hardware. These results demonstrate reliable performance across heterogeneous imaging conditions. By reducing processing time from hours of manual effort to approximately two minutes per subject, the proposed fully automated workflow facilitates routine clinical deployment and scalable multi-center studies.

## 1. Introduction

Epilepsy affects more than 50 million people worldwide and approximately one-third of patients suffer from drug-resistant epilepsy (DRE) [1, 2]. For patients with focal DRE, surgical intervention remains the most effective treatment, offering the possibility of freedom of seizures or a substantial reduction in seizure burden [3, 4]. Accurate localization of the seizure onset zone (SOZ) is therefore critical for successful surgical planning.

Stereoelectroencephalography (SEEG) is widely regarded as the gold standard for invasive localization of the epileptogenic zone in patients with drug-resistant epilepsy [5]. By stereotactically implanting depth electrodes along predefined trajectories, SEEG enables three-dimensional sampling of cortical and subcortical structures with millimeter-scale spatial resolution [6, 7]. Compared with subdural grid implantation, SEEG enables access to deep and bilateral brain structures, facilitating network-level understanding of epileptic propagation [8]. Accurate anatomical localization of each electrode contact is essential for reliable functional interpretation and connectivity analysis, which in turn inform surgical decision-making.

Traditionally, SEEG contacts localization relies on manual or semi-automated approaches, often guided by prior knowledge from surgical planning systems, such as predefined entry and target points or known trajectories [9, 10, 11]. While these prior knowledge may be available under surgical planning system, their labeling process are largely labor-intensive, operator-dependent, and limited in scalability across large datasets or multi-center studies.

Recently, fully automated pipelines have been developed to reduce manual intervention. Early rule-based methods segmented electrodes using intensity thresholding and geometric constraints [12], while volume-based localization and automated labeling strategies improved efficiency and robustness [13]. More recent end-to-end frameworks, including EpiTools [14], iEEG-recon [15], seegloc [16], and sEEG-Suite [17], integrate CT–MRI co-registration, electrode reconstruction, contact localization, and anatomical labeling into unified workflows. Despite these advances, there exist three main challenges. First, most methods rely on fixed or manually tuned intensity thresholds, which bring challenges to physicians and often fails under heterogeneous acquisition protocols or low-resolution CT conditions. Second, contact boundaries may be poorly resolved, causing contact-wise segmentation to fail when neighboring contacts appear merged. Third, electrodes may be curved or densely inserted, undermining straight-line assumptions for electrodes which are commonly adopted in these reconstruction algorithms [18].

To address these issues, we propose a fully automated and noise-robust SEEG contacts localization framework. The proposed method introduces: (1) adaptive thresholding for trajectory reconstruction without manual tuning; (2) flexible geometry-based contact localization robust to indistinct boundaries; and (3) trajectory modeling and spatial disambiguation capable of handling curved and densely implanted electrodes. We evaluate the framework on multi-center clinical data and a publicly available dataset, demonstrating recall of 98.3%, precision of 97.1% and deviation of 0.248 ± 0.066 mm for contacts localization without any manual intervention.

## 2. Materials and Methods

### 2.1. Validation datasets

The proposed pipeline was validated on two datasets with different image resolution and acquisition conditions to evaluate its robustness under practical clinical environment. One dataset consisted of five patients (Subject A to G) from Guangdong Sanjiu Brain Hospital (centen I), while an additional case with different spatial resolution was obtained from publicly available example data provided by Janca et al. (center II) [11].

For the Guangdong Sanjiu Brain Hospital, post-implantation CT (post-CT) and pre-implantation T1-weighted MRI (pre-MRI) were retrospectively collected under institutional ethics approval (Sanjiu, LSZ 2024-01-022). This dataset comprised a total of 45 SEEG electrodes with 556 contacts across five patients. The CT images in this cohort had relatively high spatial resolution (< 0.5 mm).

The publicly available dataset included post-CT and pre-MRI scans from a single patient, comprising 12 SEEG electrodes with 96 contacts, acquired at a spatial resolution of 1.0 mm. At this resolution, individual electrodes are less clearly distinguishable.

All datasets were fully anonymized prior to analysis.

### 2.2. Overview of the proposed pipeline

An overview of the proposed fully automated SEEG contacts localization pipeline is illustrated in Fig. 1. The pipeline takes post-CT and pre-MRI as inputs and outputs spatially and anatomically labeled SEEG contact coordinates in the MRI space. All processing steps are performed automatically without manual intervention.

**Figure 1:**
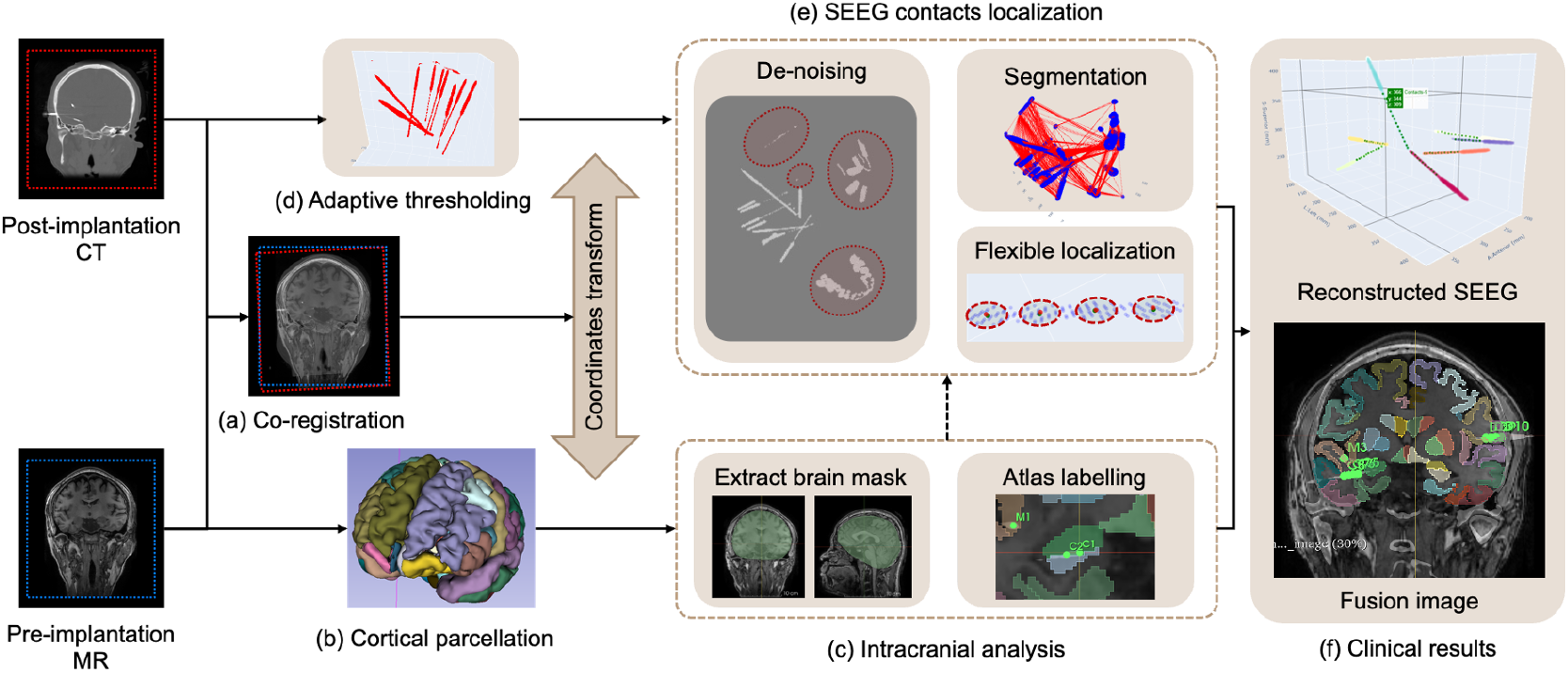
Overview of the proposed fully automated SEEG contacts localization pipeline. (a) Post-implantation CT is rigidly co-registered to the pre-implantation T1-weighted MRI to enable anatomical integration. (b) Cortical parcellation is performed on the MRI using the DKT atlas, providing anatomical context for subsequent analysis. (c) An intracranial brain mask is extracted from the cortical parcellation and used to suppress extra-cranial metallic artifacts in CT; after contact localization, detected contacts are mapped back to the parcellation to obtain anatomical labels. (d) Adaptive thresholding is applied to the CT volume to estimate an optimal threshold value for electrode segmentation under diverse imaging conditions. (e) SEEG electrode and contact localization, including de-noising using the intracranial mask, segmentation of electrode structures via spatial connectivity, and flexible contact localization by detecting intensity-density peaks along the estimated electrode principal axis, enabling robustness to lower resolution and electrode bending. (f) Reconstructed electrode trajectories and contact coordinates are transformed into MRI space and fused with anatomical parcellation results to generate spatially and anatomically labeled SEEG contacts.

### 2.3. Co-registration and cortical parcellation

Raw post-CT and pre-MRI data were initially stored in Digital Imaging and Communications in Medicine (DICOM) format. All images were converted to Neuroimaging Informatics Technology Initiative (NIfTI) format using the `dicom2nifti` Python package [19], ensuring standardized voxel representation for subsequent processing.

As shown in Fig. 1(a), rigid co-registration between post-CT and pre-MRI was performed using Advanced Normalization Tools (ANTs) [20]. The rigid transformation estimated a transformation matrix *T* mapping the post-CT space to the pre-MRI space, along with a resampled CT image, such that

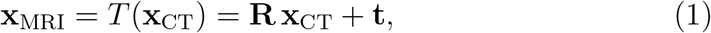

where **R** ∈ ℝ^3×3^ is an orthogonal rotation matrix (**R**^T^**R** = **I**, det(**R**) = 1) and **t** ∈ ℝ^3^ is a translation vector. The transformation matrix T allows electrode trajectories and contact coordinates to be transformed between MRI and CT spaces.

Cortical parcellation was performed using the Desikan–Killiany–Tourville (DKT) protocol [21] implemented in ANTsPyNet [22], as shown in Fig. 1(b). This procedure produced a cortical atlas comprising 101 anatomical regions. The resulting parcellation was subsequently used to generate an intracranial brain mask and to assign anatomical labels to localized SEEG contacts after spatial transformation into the MRI space.

### 2.4. Adaptive thresholding of post-implantation CT

Binary thresholding of CT is essential in preprocessing within SEEG contacts localization, as it converts volumetric post-CT data into a sparse voxel representation focusing on high-density metallic structures corresponding to implanted electrodes.

Most existing automated approaches employ a single, manually defined CT intensity threshold [23, 11, 16, 18, 14, 15], or a fixed percentile-based threshold [13]. However, in practical clinical settings, post-CT images acquired crossing different centers and scanners exhibit substantial variability in intensity distributions, making a fixed threshold unreliable. In addition, the optimal threshold values may vary from electrodes even within the same scan.

In this work, we proposed an adaptive thresholding strategy based on statistical analysis of the post-CT Hounsfield Unit (HU) intensity distribution.

In post-CT images, voxels corresponding to air and skull dominate the lower-to-moderate HU range, while SEEG electrodes occupy only a small fraction of voxels with HU values typically exceeding 2500. Examination of the HU histogram reveals that voxel counts remain high across low HU values and then sharply decline in the 1500–2500 HU range. The trailing edge of this rapid decrease reliably captures all electrode-related voxels while suppressing the majority of non-electrode structures. The optimal segmentation threshold is automatically derived from the peak of the second derivative of the HU distribution, with a slight offset applied to ensure robust extraction of the electrodes.

### 2.5. SEEG contacts localization

#### 2.5.1. De-noising of high-density non-electrode artifacts

Following adaptive thresholding, the post-CT volume may still contain high-density non-electrode structures, including dental components and metallic artifacts from extracranial electrode wires and device, as illustrated in Fig. 1(e). These structures interfere with subsequent procedures of electrode segmentation and contact localization, and must be removed prior to further processing.

To this end, a brain mask is generated from the cortical parcellation of the pre-implantation MRI by computing an convex hull of intracranial regions, as shown in Fig. 1(c). The resulting brain mask is mapped from the MRI space to the CT space using the rigid co-registration transformation and then applied to the thresholded post-CT volume to remove high-density voxels located outside the intracranial region.

#### 2.5.2. Clustering of electrode structures

After de-noising, the remaining high-density voxels predominantly correspond to implanted SEEG electrodes. These voxels are spatially organized into multiple elongated structures, each representing a single electrode trajectory, as illustrated in Fig. 1(e).

To cluster individual electrode structures and segment the densely stuck electrode, a three-dimensional connectivity analysis is performed on the denoised voxel set. Connected components are identified using depth-first search (DFS) [24] and grouped into distinct voxel clusters based on spatial adjacency. Each resulting cluster is treated as an individual electrode structure and processed independently in subsequent contact localization steps.

#### 2.5.3. Flexible contact localization along electrode trajectories

For each segmented electrode structure, the three-dimensional coordinates of its voxel cluster are analyzed to estimate the electrode trajectory. Principal component analysis (PCA) [25] is applied to the voxel coordinates to obtain the dominant axis of the electrode, which serves as a one-dimensional reference direction for contact localization.

The voxel intensities are then projected onto the estimated electrode axis, and the distribution of CT intensity along this axis is analyzed without any prior knowledge of electrode contacts. Local intensity peaks corresponding to individual electrode contacts are identified along the projected axis in a fully data-driven manner. For each detected peak, a flexible localization is performed by computing the centroid of voxels within a local neighborhood around the peak, yielding the final contact center position.

This procedure enables robust contact localization under lower resolution imaging conditions and accommodates electrode bending without assuming fixed inter-contact spacing or rigid electrode geometry, as illustrated in Fig. 1(e).

### 2.6. Electrode reconstruction and anatomical labeling

Following contact localization, the detected contact centers for each electrode are ordered along the estimated electrode trajectory to reconstruct the complete SEEG electrode geometry. Each contact is associated with its corresponding electrode shaft based on the voxel cluster identified during electrode structure segmentation.

As illustrated in Fig. 1(f), the final output of the pipeline consists of spatially localized SEEG contact coordinates in the MRI space, together with their corresponding electrode identities and anatomical labels, enabling direct integration with downstream analyses and pre-surgical evaluation workflows.

## 3. Results and Discussion

### 3.1. Automated SEEG localization pipeline

The performance of the proposed fully automated SEEG contacts localization pipeline was evaluated on datasets from six patients with varying imaging quality and acquisition conditions. All results were independently verified by a clinical expert specialized in SEEG contact localization.

For evaluation, all contacts were manually annotated by the clinical expert following the standard clinical workflow (exploiting SEEGA tools [10] with manual verification), and these annotations were regarded as the ground truth. The proposed automated pipeline was then applied to localize the contacts, and the results were compared with the ground truth by computing the Euclidean distance between corresponding contact centers. A detected contact was considered incorrect if the Euclidean distance exceeded 1 mm or if the clinical expert judged the localization to be inaccurate; otherwise, it was considered correct.

For each subject *i*, precision and recall were reported as metrics for results evaluation of the proposed automated pipeline.

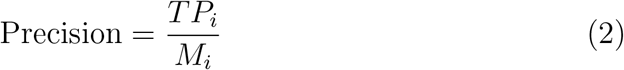

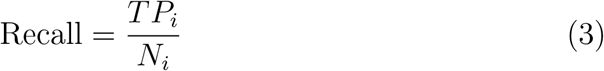

where *TP*_*i*_ represents the number of correctly localized contacts, *N*_*i*_ corresponds to the total number of the implanted contacts and *M*_*i*_ is the total number of the generated labels produced by the automated pipeline.

In addition, the mean and standard deviation of the Euclidean distances between correctly localized contacts and the ground truth were computed to quantify localization deviation.

Figure 2 presents representative qualitative results from two cases with distinct CT resolutions. Patient A from Center I (Fig. 2(a)) represents a case with post-CT in 0.5 mm resolution, in which SEEG contacts are clearly delineated. Patient F from Center II (Fig. 2(b)) represents a more challenging scenario with post-CT in 1.0 mm resolution and increased imaging noise. Despite these differences, the proposed pipeline successfully localized SEEG contacts, reconstructed electrode trajectories, and mapped the results into the pre-implantation MRI space in both cases.

**Figure 2:**
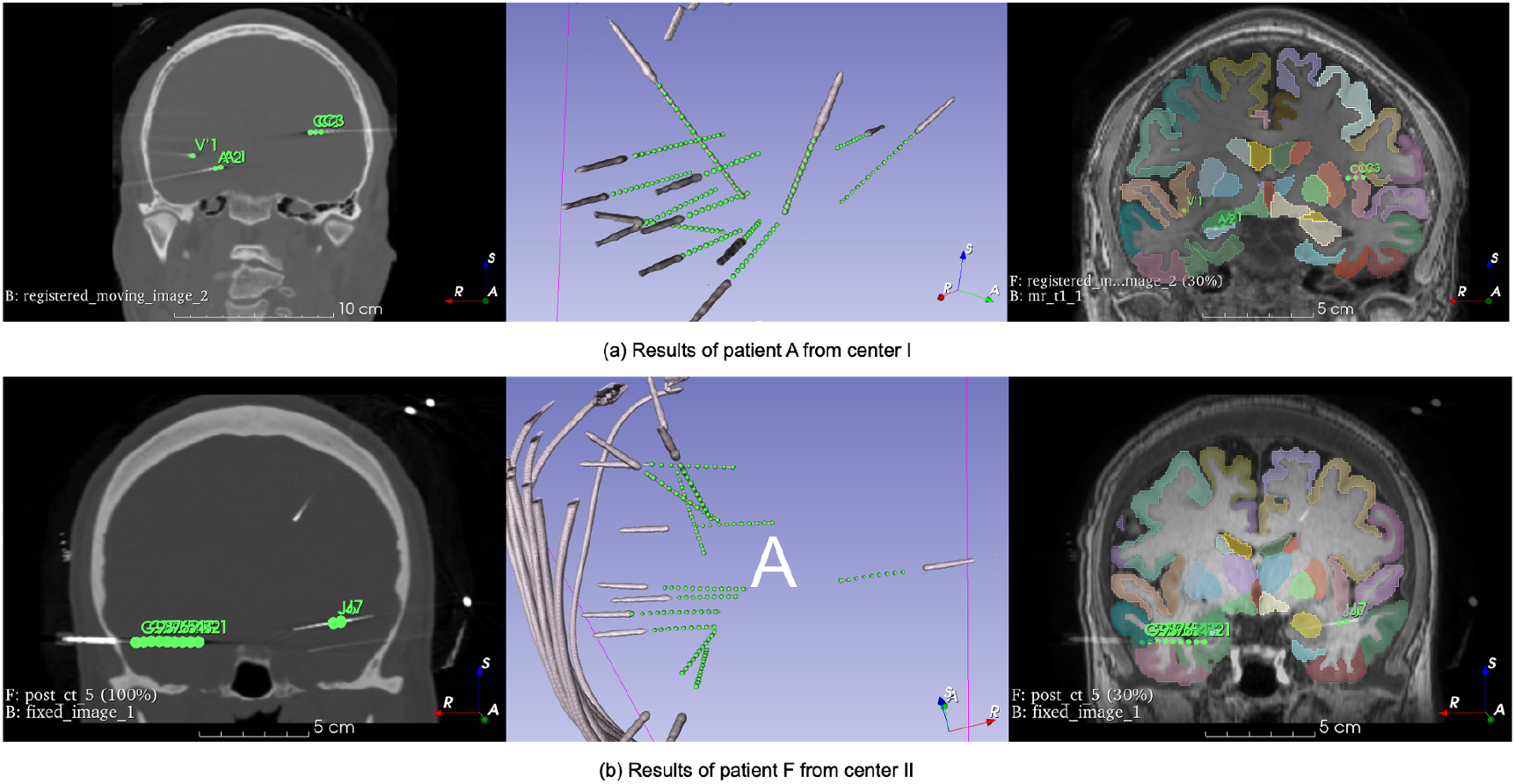
Results of the proposed fully automated SEEG contacts localization pipeline. Results of patient A from center I with 0.5 mm imaging resolution. (b) Results of patient F from center II with 1.0 mm imaging resolution. For each case, the left panel shows SEEG contacts localized in the post-implantation CT space after adaptive thresholding and denoising; the middle panel shows the reconstructed electrode trajectories with spatially ordered contact sequences; the right panel shows the final multimodal fusion visualization, integrating CT, pre-implantation MRI, cortical parcellation, and localized SEEG contacts. All visualizations were rendered in 3D Slicer software.

As shown in Fig. 2, localized contacts form spatially ordered sequences along reconstructed electrode trajectories and are anatomically consistent with intracranial structures. The final multimodal fusion visualization enables direct inspection of each SEEG contact within its corresponding cortical parcellation region. All results were obtained automatically without manual parameter tuning or user interaction.

Results of quantitative evaluation are summarized in Table 1. Across all six patients, the proposed automated pipeline successfully reconstructed all implanted electrodes and achieved an overall recall of 98.3% and precision of 97.1% for contact localization. The small proportion of missed contacts primarily occurred in regions with densely clustered electrodes, where minor segmentation imperfections generated spurious artifacts that interfered with accurate contact detection.

**Table 1:** Performance of the automated SEEG contacts localization pipeline.

| Patient | Center | Electrodes | Contacts ($N_i$) | Predicted ($M_i$) | True ($TP_i$) | Precision | Recall |
| --- | --- | --- | --- | --- | --- | --- | --- |
| A | I | 10 | 129 | 131 | 129 | 0.985 | 1.000 |
| B | I | 7 | 90 | 90 | 87 | 0.967 | 0.967 |
| C | I | 13 | 168 | 169 | 166 | 0.982 | 0.988 |
| D | I | 9 | 108 | 110 | 105 | 0.955 | 0.972 |
| E | I | 6 | 61 | 63 | 59 | 0.937 | 0.967 |
| F | II | 12 | 96 | 97 | 95 | 0.979 | 0.990 |
| Total | – | 57 | 652 | 660 | 641 | **0.971** | **0.983** |

The deviation of correctly localized contacts is presented in Table 2. The overall localization deviation was 0.248 ± 0.066 mm, indicating high spatial consistency with the expert-annotated ground truth.

**Table 2:** Deviation of correctly localized SEEG contacts.

| Patient | Mean Error (mm) | Std (mm) |
| --- | --- | --- |
| A | 0.157 | 0.041 |
| B | 0.425 | 0.092 |
| C | 0.193 | 0.058 |
| D | 0.288 | 0.076 |
| E | 0.392 | 0.085 |
| F | 0.171 | 0.049 |
| Overall | **0.248** | **0.066** |

These results demonstrate the robustness and high precision of the proposed fully automated localization pipeline across datasets with varying image quality.

### 3.2. Methodological validation

The key innovations of the proposed SEEG contacts localization pipeline were systematically validated, as illustrated in Fig. 3.

**Figure 3:**
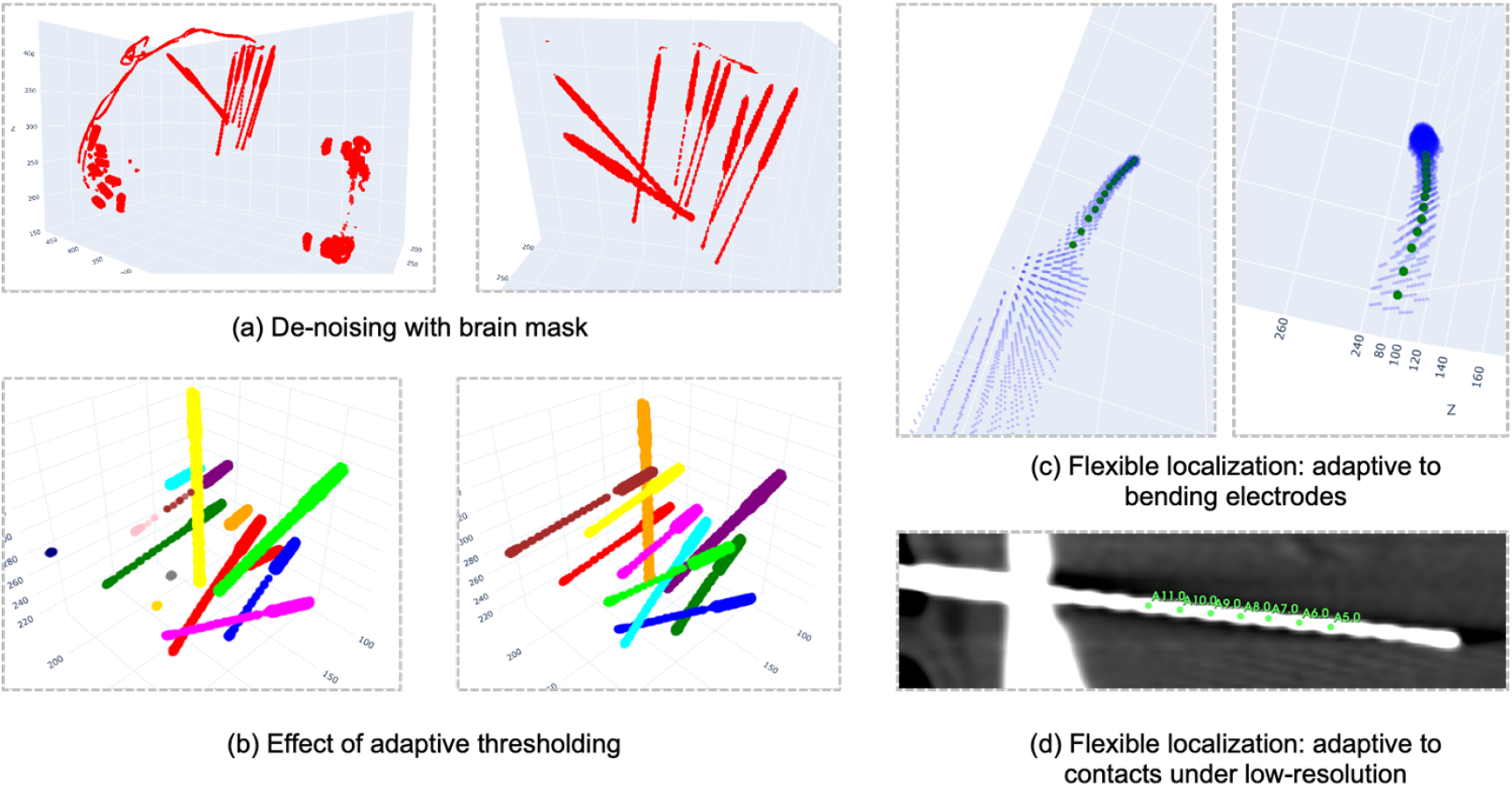
Validation of key methodological components of the proposed automated SEEG contacts localization pipeline. (a) Denoising effect using brain mask: raw volume (left) and de-noised volume (right). (b) Adaptive thresholding: comparison between fixed (left) and adaptive (right) thresholds. (c) Flexible contact localization for bended electrodes. (d) Flexible contact localization under lower resolution conditions.

Figure 3(a) demonstrates the effectiveness of the denoising step. By applying a brain mask, most extracranial artifacts, including residual skull fragments and metal noise, are removed, preventing false detections of electrodes and improving downstream localization reliability.

The adaptive thresholding strategy was evaluated in Fig. 3b. When using a fixed threshold (left panel), some electrodes are only partially visible due to CT acquisition angle or dose limitations, showing only the bolt and sparse contact targets. Conventional automated methods fail to properly localize these electrodes. By contrast, the adaptive thresholding approach (right panel) accurately reconstructs all electrodes for each case, enhancing the sensitivity and completeness of the automated localization.

Figures 3(c) and 3(d) illustrate the performance of the flexible contact localization strategy. For curved electrodes (Fig. 3(c)), the method accurately captures the full electrode trajectory and adapts to bending, outperforming traditional straight-line assumptions in terms of robustness and precision. Under low-resolution conditions (Fig. 3(d)), where contacts may appear connected or partially merged, the method still reliably localizes contacts by identifying local HU peaks corresponding to the contact centers. This demonstrates strong anti-interference capability and maintains localization accuracy even in challenging imaging scenarios.

Overall, these validation results confirm that the proposed methodological innovations—denoising, adaptive thresholding, and flexible contact localization—significantly improve the robustness, sensitivity, and accuracy of fully automated SEEG contacts localization across diverse imaging conditions.

### 3.3. Computational efficiency

The computational efficiency of the proposed fully automated SEEG contacts localization pipeline was evaluated on a standard desktop workstation equipped with an Intel Core i7-1260P 2.10 GHz processor and 32 GB of RAM. Pipeline processing times were recorded for all six patient datasets, and the results are summarized as mean ± standard deviation.

On average, co-registration of post-implantation CT to pre-implantation MRI required 35.7 ± 5.1 s per case. The cortical parcellation and brain masking steps took 58.9 ± 4.3 s. Automated SEEG electrode and contact localization was completed in 20.6 ± 2.5 s. Overall, the total pipeline execution time was 120.2 ± 8.6 s per case.

These results indicate that the proposed pipeline can process a full SEEG dataset in approximately two minutes on standard computing hardware, while traditional mannual operation costs up to 1 hour to acquire the same tasks. This demonstrates our potential suitability for routine clinical use and high-throughput research applications.

### 3.4. Comparison with other approaches

We further compared the proposed pipeline with two existing SEEG localization tools, VoxTool [15] and EpiTools [14]. Precision and recall were computed for each patient, and overall performance was summarized across all cases, as shown in Table 3.

**Table 3:**
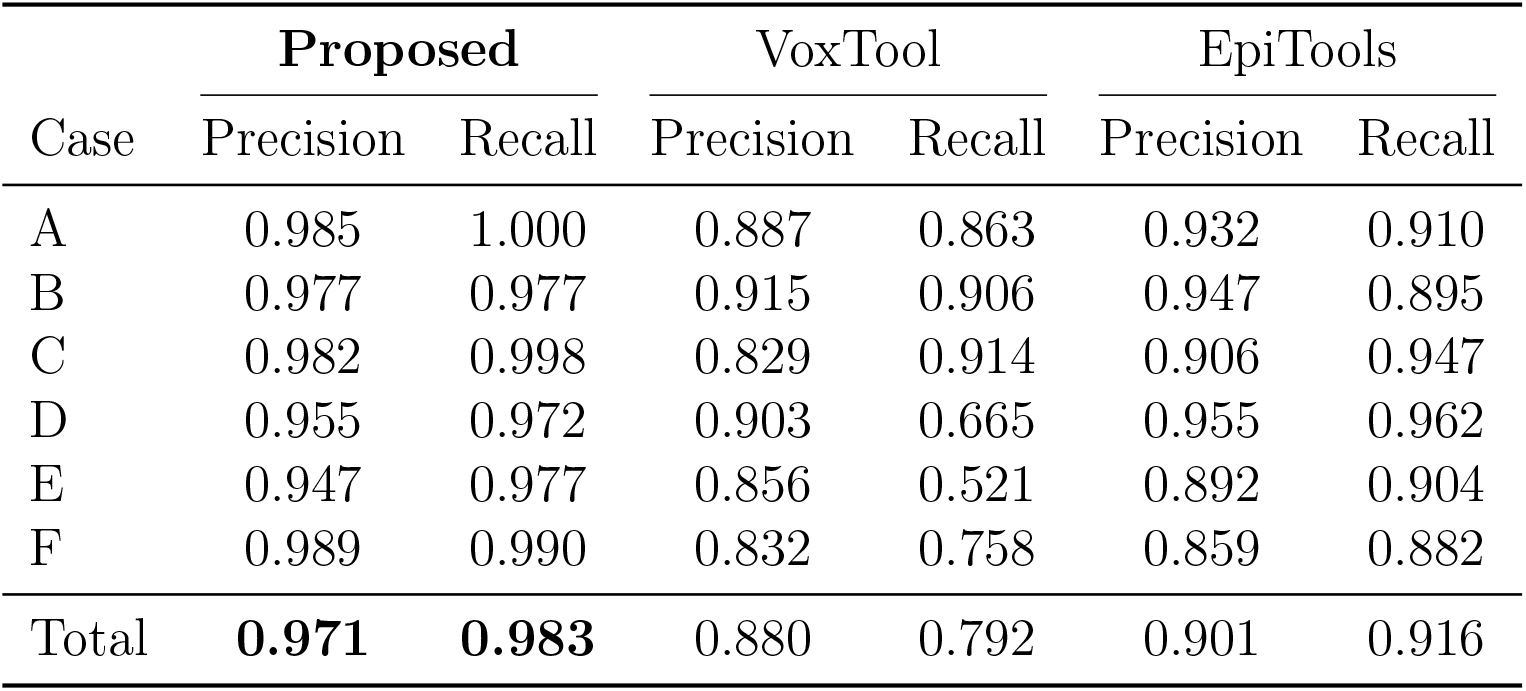
Comparison of precision and recall with other approaches.

Overall, the proposed pipeline achieved a precision of 0.971 and a recall of 0.983, outperforming VoxTool (precision 0.880, recall 0.792) and EpiTools (precision 0.901, recall 0.916). Notably, the proposed method maintained consistently high recall across all patients, indicating robust localization of SEEG contacts under heterogeneous imaging conditions.

These findings indicate that the proposed framework achieves competitive and consistently high performance across subjects, with improved recall compared to existing approaches.

## 4. Conclusion

In this study, we present a fully automated SEEG contacts localization pipeline that requires no manual intervention or parameter tuning and is adaptable to multi-center datasets. The pipeline integrates adaptive thresholding to extract optimal binary thresholds in various CT conditions, reconstructing complete electrode structures, and a flexible contact localization strategy to handle bent electrodes and low-resolution imaging. Preliminary validation on datasets from two centers, encompassing various resolution images, demonstrates that our method produces robust and reliable localization results with high computational efficiency. These features make the pipeline well-suited for clinical deployment and scalable studies, providing a solid foundation for future large-scale data validation and integration into presurgical workflows.

## Acknowledgment

The authors would like to thank the clinical teams at Guangdong Sanjiu Brain Hospital and the contributors of the publicly available SEEG dataset for providing access to patient imaging data.

